# Two Forms of Knowledge Representations in the Human Brain

**DOI:** 10.1101/691931

**Authors:** Xiaoying Wang, Weiwei Men, Jiahong Gao, Alfonso Caramazza, Yanchao Bi

**Author notes:** Correspondence to Y. Bi.

## Abstract

Sensory experience shapes what and how knowledge is stored in the brain -- our knowledge about the color of roses depends in part on the activity of color-responsive neurons based on experiences of seeing roses. We study the brain basis of color knowledge in congenitally blind individuals whose color knowledge can only be obtained through language descriptions. We found that some regions support color knowledge only in the sighted. More importantly, a region in the left dorsal anterior temporal lobe supports object color knowledge in both the blind and sighted groups, indicating the existence of a sensory-independent knowledge coding system in both groups. Thus, there are (at least) two forms of object knowledge representations in the human brain: sensory-derived and cognitively-derived knowledge, supported by different brain systems.

## Introduction

Roses are red, violets are blue, so goes the poem. What is the neural code for such knowledge in the human brain? Although object knowledge can be acquired both through sensory experience (seeing red roses) and through language description (e.g., being told by others that roses are “red”), one prominent theory assumes that such knowledge is stored in sensory association cortices, derived from sensory experiences (i.e., seeing the colors of roses) (Barsalou et al., 2003; Martin, 2016; Simmons et al., 2007). The alternative possibility is that even sensory-derived knowledge is also represented at an abstract conceptual level distinct from sensory representations (Leshinskaya and Caramazza, 2016; Mahon and Caramazza, 2008; Shallice, 1988, 1987; Wang et al., 2018; Xu et al., 2017).

Prior attempts to test the existence of and to characterize the neural representation of non-sensory derived object knowledge include studies of congenitally, sensory-deprived populations (e.g., blind/deaf) and studies, with sighted and blind individuals, of abstract concepts such as “justice” or “idea”, for example, that do not refer to specific sensory experiences. However, it has been argued that even such concepts may still be grounded in their associated sensory, emotional and complex event experiences (Barsalou, 2008; Borghi and Binkofski, 2014; Kiefer and Pulvermüller, 2012; Kousta et al., 2011; Vigliocco et al., 2014). And, in the case of studies based on sensory deprivation, aspects of the knowledge not available from one sense could be obtained partially through other senses (Amedi et al., 2007; Ricciardi et al., 2013). For instance, object shape knowledge can be obtained both by vision and by touch (Lacey and Sathian, 2014, 2012; Peelen et al., 2014). Given these considerations, color knowledge in congenitally blind individuals is especially appropriate for the investigation of conceptual representations. Congenitally blind people, without any way to sense color, respond similarly as sighted people to color-related questions(Connolly et al., 2007; Landau, 1983; Landau and Gleitman, 1985; Marmor, 1978; Saysani et al., 2018; Shepard and Cooper, 1992). In these individuals, color knowledge can only be obtained through language descriptions (e.g., some roses are red; red is more similar to orange than to blue), providing a unique opportunity to test how and where such knowledge is represented in the brain. It also allows investigation of the relationship between language-derived knowledge of object color and the corresponding sensory-derived knowledge in the typically-developed brain. By comparing the brain basis of object color knowledge in blind and sighted individuals, we found that some regions support color knowledge only in the sighted and, more importantly, we also found a region that represents object color knowledge in both groups, supporting the existence of a sensory-independent knowledge coding system, That is, object color knowledge in the typically developed brain is distributed over distinct regions that represent sensory-based and abstract conceptual information, respectively.

## Results

We compared congenitally blind (CB; see Table S1 for detailed characteristics) and sighted control (SC) individuals on their knowledge about colors of common fruits and vegetables using a set of behavioral tasks. We also compared the neural representation of object color knowledge in the two groups through fMRI experiments. In the main behavioral task, we asked subjects to judge the color similarity of pairs of objects on a 1-7 scale (Fig. 1A). Other tasks included explicitly generating color names for a given fruit or vegetable (Fig. S1, materials and methods). The behavioral results showed that the object color knowledge similarity space of the CBs, who never experienced colors, was highly similar to that of SCs. Fig. 1A shows the representational similarity matrices (RSM) obtained from the pairwise, rating-derived, object color similarities averaged across participants from each group. The group-mean color RSMs of the CB and the SC participants were highly correlated (Fig. 1A; Pearson r = 0.88, n=276, P < 3.91×10-90). The same pattern of results was also obtained using the behavioral task requiring explicit color name generation for objects (Fig. S1A). Multi-dimensional scaling (MDS)(Busing et al., 1997; Takane et al., 1977), using individual differences scaling solutions (INDSCAL), was carried out to visualize the object color space in each group. This analysis revealed two highly similar two-dimensional object spaces for the CB and SC groups (Fig. 1B; r = 0.88, P < 5.85×10-90). Within each object space, objects loosely clustered into those with red, yellow, green and purple colors, forming a spherical spectral transition of colors. A subject space was computed for each group (Fig. 1C), within which each individual participant was located according to their weights on each dimension of the object space. Comparing the dimensional weights of the two groups using mixed effects ANOVA (between-subject factor, group; within-subject factor, dimension) revealed that the two groups had similar object color space structures (indicated by insignificant interaction, F(1,25) = 0.09, P = 0.772), but the CB participants showed slightly greater individual variations on the object color space (indicated by a significant group main effect, F(1,25) = 4.63, P = 0.041; CB vs. SC, mean difference±SE = −0.003±0.002). Comparison of the within-group individual subject correlations further confirmed larger individual variability in the CB relative to the SC group (Fig. 1D; CB vs. SC: mean difference ± SE = −0.19±0.07, t(25) = −2.54, P < 0.018, two-tailed).

**Fig. 1.**
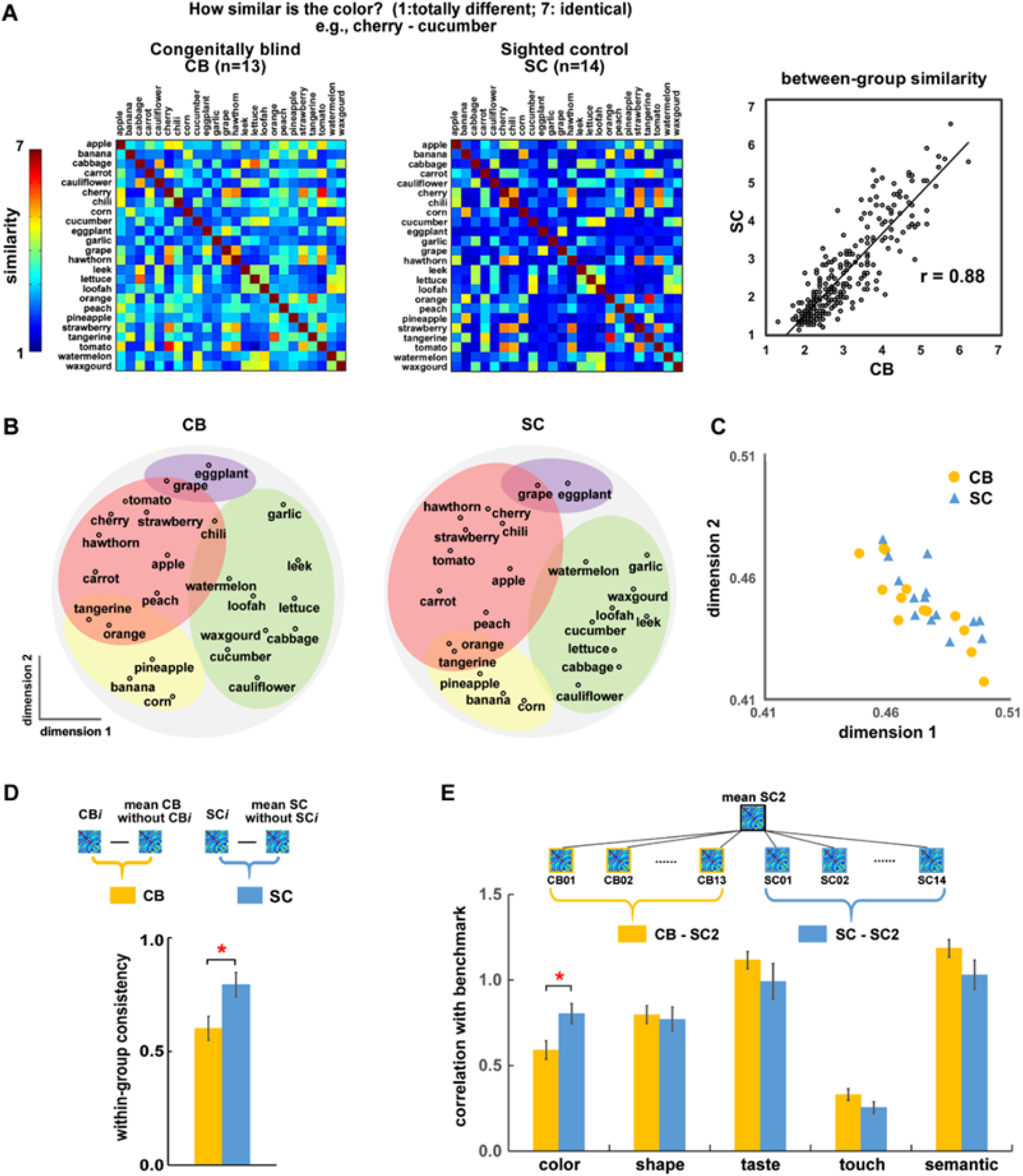
Object color representation in CB and SC participants. (A) Object color RSMs derived from pairwise object color similarity rating for CB and SC groups and the correlation between them. Color bar indicates the rating score averaged across participants in each group. (B) Object color spaces obtained through INDSCAL MDS on individual object color RSMs from each group. (C) One subject space combining the subject spaces of both CB and SC. The coordinates of each point in the space correspond to each individual subject’s weights along dimension one and dimension two (see methods and materials). (D) Correlations between each individual’s object color RSM and the corresponding leave-one-out group mean RSM for CB (yellow bar) and SC (blue bar). (E) The similarity of object color representation between CB (yellow bar) or SC (blue bar) individuals and the SC2 benchmark across different object features. Red asterisks indicate significant between-group difference. Color patches in (B) were added subjectively to highlight the clustering pattern of objects with different colors.

Taking the group-mean object color RSM of an independent group of sighted college students (SC2) as a benchmark and correlating the RSM of each CB and SC individual with it, we found significantly lower CB-SC2 correlations (Fisher-transformed r: mean = 0.59, SE = 0.05) relative to SC-SC2 correlations (Fisher-transformed r: mean = 0.80, SE = 0.06) using two-sample t-test (Fig. 1E, t(25) = −2.69, P < 0.013, two-tailed). That is, although the CB and SC individuals have largely similar object color knowledge spaces, there are also subtle yet significant differences, in line with previous findings (Marmor, 1978; Saysani et al., 2018; Shepard and Cooper, 1992). Differences between the CB and SC groups were not observed for object knowledge that could be acquired through nonvisual sensory modalities (i.e., shape, taste, touch; CB-SC2 vs. SC-SC2: ts < 1.59, Ps > 0.125). The relationship between object color knowledge and knowledge about other types of object features (i.e., shape, taste, touch) were significant in both the CB and SC groups (Fig. S2, Ps < 3.05×10-5 for both groups), with no significant between-group differences (ts < 1.68, Ps > 0.107).

The central question investigated here is whether object color knowledge, as reflected in the semantic spaces obtained for the CB and SC groups, is supported by fundamentally different neural substrates in the two groups, as might be expected from theories which assume that color knowledge representations in the sighted are based on sensory experience. FMRI scanning was conducted to address this question (Fig. 2A). Fifteen CB and twenty SC participants listened to fruit and vegetable names and judged whether the color of each fruit or vegetable is red/reddish in the real world. Comparable response profiles for the two groups of participants were obtained with this task (Fig. 2A). Before exploring the pattern of neural responses across the whole-brain, we first focused on two brain regions which have been proposed as candidates for sensory versus non-sensory abstract knowledge representation: visual color perception areas (Martin, 2016; Simmons et al., 2007) and left anterior temporal lobe (ATL) (Lambon Ralph et al., 2017; Noppeney and Price, 2004; Striem-Amit et al., 2018; Wang et al., 2010).

**Fig. 2.**
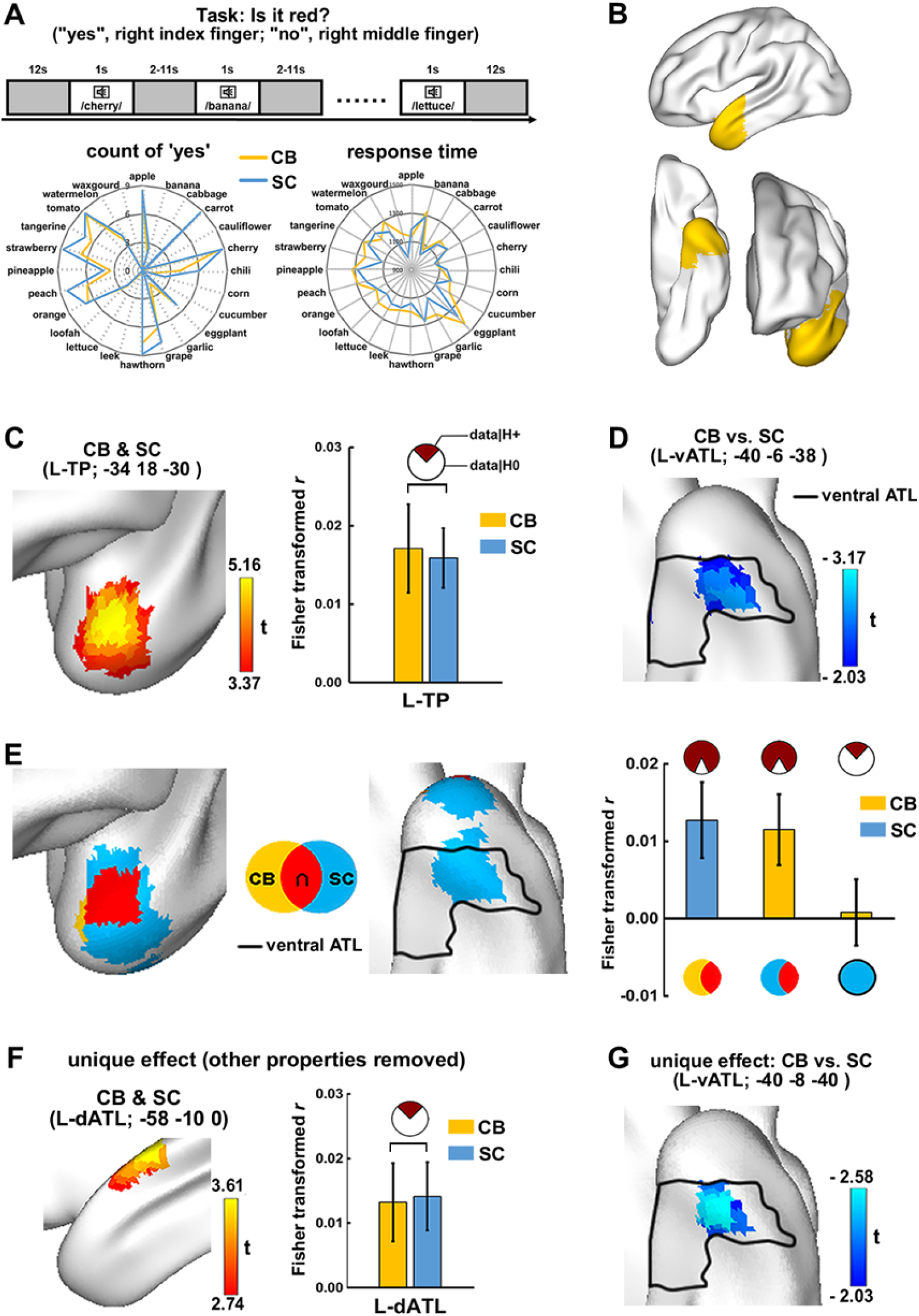
Object color representation within the left ATL. (A) Design and behavioral performances of the object color verification fMRI experiment. (B) The anatomically defined left ATL (see materials and methods in supplementary material). (C) Combination of object color effect across the CB (n=15) and the SC (n=20) groups. Voxel-level p < 0.001, one-tailed, cluster-level FWE corrected P < 0.05. The bar plot shows the magnitude of the object color effect (RSA-derived Fisher-transform r values averaged across participants) in CB and SC groups in the left TP. The pie chart depicts the results of the Bayesian t-tests, including the odds of the data under H0 (white part) vs. H1 (dark red part). (D) Between-group comparison of object color effect. Cold color indicates weaker effect in CB relative to in SC. Voxel-level P < 0.05, two-tailed, 33 voxels, uncorrected. (E) The object color effect in CB (yellow) and SC (blue) respectively and their overlap (red). The solid line encompasses the ventromedial ATL corresponding to the anterior inferior temporal gyrus, the anterior temporal fusiform cortex and the anterior parahippocampal gyrus in the anatomically defined left ATL. Voxel-level P < 0.01, one-tailed, cluster-level FWE corrected P < 0.05 in each group. The bar plot shows the cross-group ROI analysis results: The object color effect of SC group (blue bar) in CB’s significant object color cluster (red and yellow patches in E) and the object color effect of CB group (yellow bars) in SC’s significant object color cluster within the ventromedial ATL (the blue patch circled by solid line in E) and outside it (the red and blue patch outside the solid line in E). (F) Combination of unique object color effect across CB and SC groups. The effects of object shape, taste, touch and semantic similarity were partialled out. (G) Between-group comparison of unique object color effect. Voxel-level P < 0.05, two-tailed, 27 voxels, uncorrected.

In the anatomically defined left ATL (Fig. 2B), representation similarity analysis (RSA) searchlight mapping (Kriegeskorte, 2008; Kriegeskorte et al., 2006), which examines the relationship between the regional neural activity pattern and object color knowledge RSMs (Fig.1A), revealed two clusters with different profiles (Fig. 2C–2E): a dorsal anterior ATL cluster where both SC and CB showed effects of object color knowledge representation; and a ventral ATL cluster where only SC showed significant effects of object color knowledge representation. Specifically, SnPM-based one sample t-test on the RSA searchlight mapping results combining the two groups revealed a significant cluster in the left dorsal temporal pole (L-TP; Fig. 2C, voxel-level P < 0.001, one-tailed, cluster-level FWE corrected P < 0.05), with no difference between the two groups in the ROI-based RSA (Fig. 2C, bar plot). Analyses of the object color knowledge representation in each group separately converged on finding this overlapping dorsal ATL cluster (Fig. 2E, red patch; for each group SnPM-based one-sample t-test, voxel-level P < 0.01, one-tailed, cluster-level FWE corrected P < 0.05). The significant cluster in the sighted group further extended to the ventral ATL (Fig. 2E, blue patch circled by solid line). Bayesian one-sample t-tests (H1: group mean > 0) on the ROI-based RSA results revealed that the SC group’s object color knowledge effect was significant (Fig 2E bar plot, blue bar, BF10 = 6.33, moderate evidence in favor of H1) in the CB group’s significant cluster (the red and yellow patch in Fig. 2E); The CB group’s object color knowledge effect was also significant (Fig. 2E bar plot, yellow bar in the middle, BF10 = 3.55, moderate evidence in favor of H1) in the anterior dorsal part of the SC group’s significant cluster (the red and blue patch outside the solid line in Fig. 2E). No evidence was observed for object color knowledge representation in the CB group within the ventral part of the SC group’s significant cluster (the blue patch enclosed by the solid line in Fig. 2E; Fig. 2E bar plot, yellow bar at right, BF10 = 0.30, moderate evidence in favor of H0). The ventral part of ATL also showed trends of stronger object color knowledge effect in the SC group relative to the CB group (voxel-level P < 0.05, two-tailed, cluster size = 34 voxels, uncorrected; Fig. 2D).

The object color knowledge representation effects in ATL are not attributable to other object features that correlated with object color knowledge. After controlling for the effects of object shape, taste, touch and general semantics using partial correlation including the RSMs of these properties as covariates (RSMs separately obtained by asking subjects to rate the similarities of object pairs on each of these features; Fig. S2), there was still a dorsal cluster showing a significant unique object color effect that was a bit posterior and dorsal to the peak obtained without controlling covariates (combining the two groups, Fig. 2F; voxel-level P < 0.005, one-tailed, cluster-level FWE corrected P < 0.05). The peak cluster before controlling for the covariates still showed trends of effects in the left TP: voxel-level P < 0.005, one-tailed, k=39, uncorrected, not shown in the figure. Comparison between the CB and SC groups in the left ATL revealed trends of stronger unique object color representation in SC relative to CB participants in the ventral ATL (Fig. 2G; voxel-level p < 0.05, two-tailed, k = 27, uncorrected). The results also held when using the behavioral results of the other object color knowledge tasks for the RSA (Fig. S3).

Within the color perception mask, which was based on a color perceptual functional localizer (Beauchamp, 1999; Simmons et al., 2007) collected from a group of 14 sighted participants (Fig. 3A), clusters showing unique object color knowledge representation wa observed only for the SC group, in the posterior parts of the right inferior temporal gyrus (ITG) and the left posterior fusiform gyrus (pFG) (Fig. 3B; voxel-level P < 0.001, one-tailed, cluster size > 10 voxels, uncorrected). The CB group showed no effect of unique object color knowledge representation in either ROI-based RSA in these two clusters (Fig. 3B bar plot; Bayesian on sample t-test, H1: group mean > 0; L-pFG, BF10 = 0.10, strong evidence in favor of H0; R-ITG, BF10 = 0.16, moderate in favor of H0) or in the RSA searchlight mapping (no cluster shown at uncorrected P < 0.01, one-tailed, cluster size > 10 voxels). Between-group comparison revealed significantly stronger object color knowledge representation in the SC relative to the CB group in the left pFG (Fig. 3C; voxel-level P < 0.005, two-tailed, cluster-level FWE corrected P < 0.05). Similar patterns of results were observed for object color knowledge effect without controlling for other object features (Fig. S4).

**Fig. 3.**
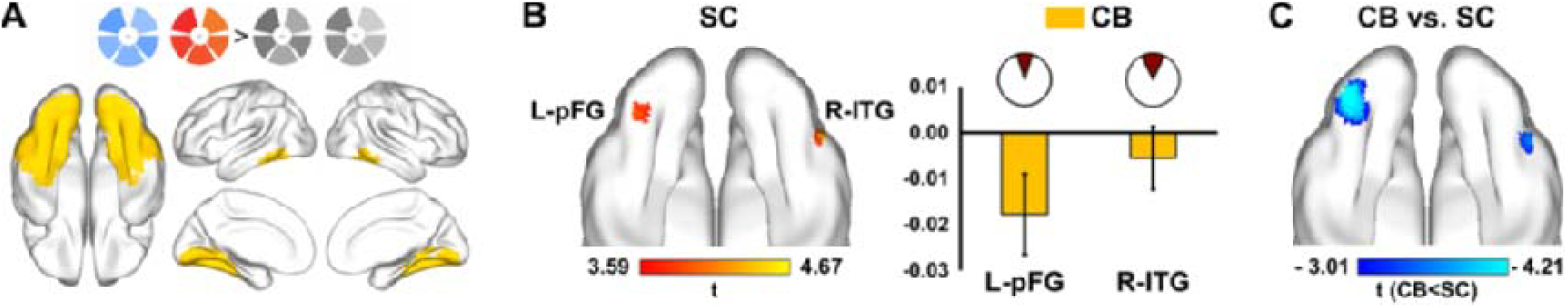
Object color representation within the color-sensitive VOTC. (A) Color-sensitive VOTC regions defined from the color perceptual localizer (see materials and methods). (B) Regions showing the unique object color effect in the SC group (voxel-level P < 0.001, one-tailed, cluster size > 10 voxels, uncorrected). The bar plot shows the CB’s unique object color effect in these regions. (C) Between-group comparison of the unique object color effect.

The intrinsic resting-state functional connectivity (rsFC) profile of the three object color knowledge representation nodes obtained above — the vision-independent left anterior dorsal ATL (L-adATL; combining TP in Fig. 2C & dATL in Fig. 2F), the vision-dependent ventral ATL (L-vATL; Fig. 2G), and the posterior ventral occipitotemporal color perception node (pVOTC; combining the left pFG and right ITG in Fig. 3C) — showed interesting differences between the CB and the SC participants (Fig. 4A; ROI-wised connectivity, Bonferroni corrected P < 0.05, number of multiple comparisons = 9). While for the CB group there was no significant connection between vision-independent and vision-dependent color nodes, for the SC group these seeds are tightly connected. The language system approximated by the contrast between sentences and nonword lists (Fedorenko et al., 2010) and the color perceptual system (color-sensitive regions defined from our functional localizer; see materials and methods) were further included in the network as theoretically driven target networks. For the CB participants the vision-independent L-adATL cluster was significantly connected with the language system, and the vision-dependent pVOTC with the color perceptual system, but there was no connection between the two. For the SC participants, however, the L-adATL has connections across both of these two large systems.

**Fig. 4.**
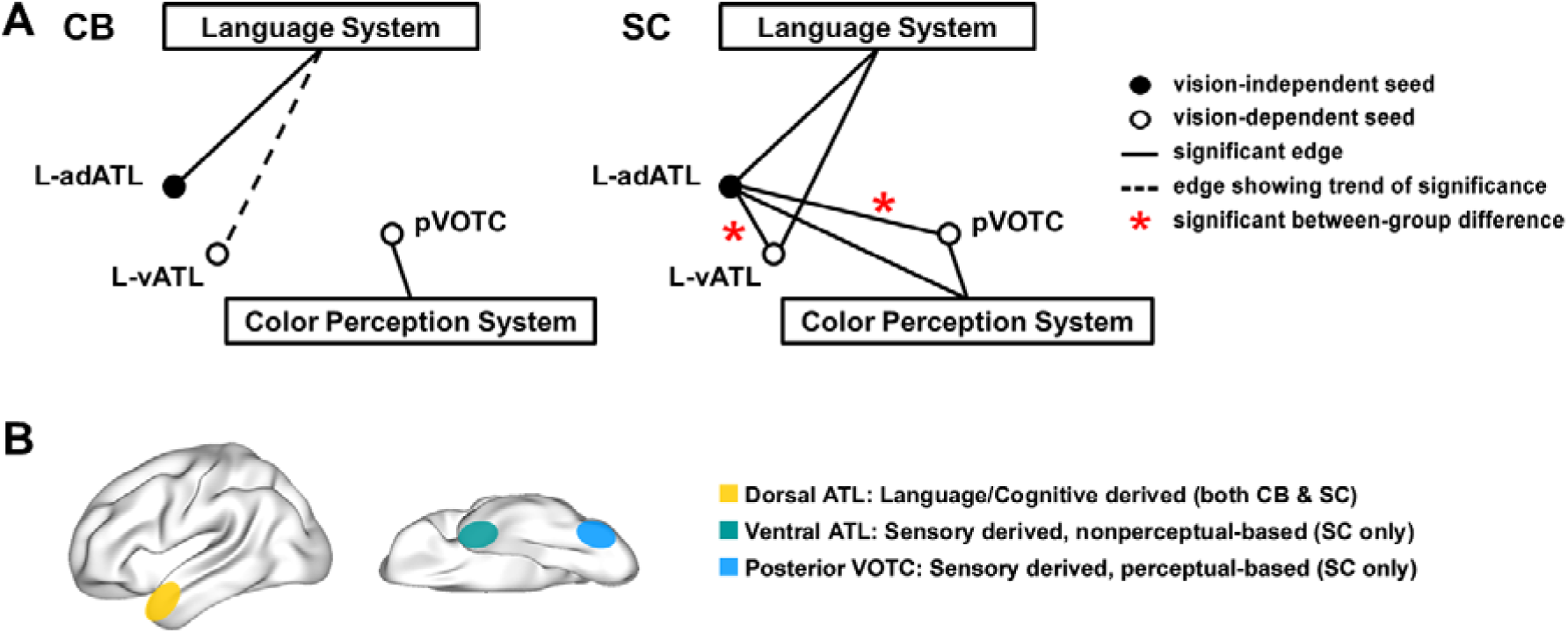
Different functional connectivity patterns in CB and SC and a schematic view of the dual forms of knowledge representation in the human brain. (A) Different rsFC networks in CB and SC group. Solid dot indicates vision-independent object color node and circles indicate vision-dependent object color nodes. Solid and dashed lines indicate significant positive connections after and before Bonferroni correction (number of comparisons = 9), respectively. Red asterisks indicate stronger connection strengths in SC relative to in CB group. (B) A schematic view of coexisting language/cognitive-derived and sensory-derived knowledge representations in the human brain. Note that there are (at least) two types of sensory-derived representations, one more likely to be based on perceptual formats (within the color perceptual system) and one more abstract (outside the color perceptual system, ventral ATL). The exact representation formats of this latter patch remain to be understood.

The object color knowledge representation results in whole-brain analyses converged well with the above results within the two pre-defined masks. Combining the two groups, RSA searchlight mapping in the whole brain revealed that clusters in the left dorsal TP and the anterior superior temporal gyrus (STG) showed significant object color knowledge representation (voxel P < 0.001, cluster-level FWE corrected P < 0.05; Fig. S5, A&C). Contrasting the two groups revealed significantly stronger effects of object color knowledge representation in the SC relative to the CB group in the posterior VOTC regions, and small clusters in the left temporoparietal, the left anterior inferior temporal and the ventral medial prefrontal cortex (Fig. S5, B&D).

Further RSA were conducted in the CB group to rule out the potential influence of the large individual variability on object color knowledge in these participants (Fig S6). We conducted RSA by excluding three CB individuals whose rating-derived object color knowledge spaces were most dissimilar from the SC group. RSA searchlight in the left ATL and ROI-based RSA results in the above-defined visual object color ROIs confirmed the results obtained with the full set of subjects: a visual-independent object color knowledge representation was obtained in the left dorsal TP (i.e., effects in both CB and SC); the effects in vATL and pVOTC regions were not observed in the CB group (Fig. S6, A-D).

Taken together, we showed that CB individuals, without any sensory experience for color, acquire object color knowledge representations highly similar to the sighted. These similar behavioral profiles in the two groups are supported by overlapping neural substrates in dorsal ATL, with the sighted participants additionally having representations in ventral ATL and VOTC color-perceptual regions. These overlaps and divergences indicate that there are two distinct formats of knowledge representation in a typically developed human brain, even for sensory-related properties (Fig. 4B): one based on sensory-derived codes (seeing the colors of roses) and one based on language/cognitive-derived codes.

## Materials and Methods

### Participants

15 congenitally blind (CB, 3 females) and 20 sighted controls (SC, 6 females) participated in the object color verification fMRI experiment. Ages and years of education were matched between CB (age: mean ± SD = 44±12, range = 24 - 65; year of education: mean ± SD = 11 ± 3, range = 0 – 15) and SC (age: mean± SD = 43 ± 11, range = 23 – 62, |t_(33)_| < 0.31; years of education: mean ± SD = 12 ± 4, range = 9 −19, |t_(33)_| < 1. 24).

13 CB (3 females) and 14 SC (4 females) with matched ages (CB: mean ± SD = 44±11, range = 22 - 63; SC: mean± SD = 40 ± 12, range = 22 - 60; t_(25)_ < 1) and years of education (CB: mean ± SD = 12 ± 2, range = 9 - 15; SC: mean ± SD = 12 ± 3, range = 9 −18; t_(25)_ < 1) participated in the behavioral tests. All of the CB and 7 of the SC participants also took part in the above object color verification fMRI experiment. The 14 SC participants also underwent a color perceptual localizer scanning.

14 CB (4 females) and 23 SC (8 females) matched in age (CB: mean ± SD = 44±11, range = 22 - 63; SC: mean± SD = 41 ± 14, range = 22 - 66; t_(34)_ < 1) and years of education (CB: mean ± SD = 12 ± 2, range = 9 - 15; SC: mean ± SD = 12 ± 4, range = 6 −18; t_(34)_ < 1) participated in an 8-minute-long (240 volumes) resting-state fMRI scan. Thirteen of the CB and 7 of the SC participants also took part in the object color verification fMRI experiment. For these participants, the resting-state scan was performed about 2 years before the object color verification fMRI experiment.

All blind participants reported that they had been blind since birth. Because medical records of onset of blindness were not available for most participants, it cannot be ruled out that some of the participants may have had vision very early in life. None of the participants remembered to have ever been able to visually recognize shapes. Five blind participants reported to have had faint light perception in the past. Three blind participants had faint light perception at the time of testing. Detailed diagnoses for all blind participants are listed in Table S1.

All participants were native Mandarin Chinese speakers. None had a history of psychiatric or neurological disorders or suffered from head injuries. All sighted participants had normal or corrected-to-normal vision and intact color perception. All participants provided informed consent and received monetary compensation for their participation. The study was approved by the Human Subject Review Committee at Peking University, China, in accordance with the Declaration of Helsinki.

### Stimuli

Twenty-four fruits and vegetables were chosen as stimuli based on high familiarity (mean ± SD = 6.13 ± 0.35, obtained from a 7-point familiarity rating on an independent group of 59 college students). The names of these fruits and vegetables were all disyllabic words with the exception of the name of carrot being tri-syllabic. For the fMRI experiment, the names were digitally recorded (22050 Hz, 16 Bit), spoken by a female native Mandarin speaker. Stimulus presentation was controlled by E-prime (Schneider et al., 2002).

### Behavioral tests

Various behavioral tests were conducted to characterize the object color representation in sighted and congenitally blind participants, which include: 1) Pairwise object feature similarity rating: 7-point similarity ratings on different object features (i.e., color, shape, taste, touch, semantic) and 2) Verbal-mediated object color knowledge and color concept tests: object color name generation and pairwise color name similarity rating (i.e., red, orange, yellow, green, cyan, blue, purple, pink, gold).

#### Pairwise object feature similarity ratings

Thirteen congenitally blind participants and 14 sighted controls matched in age and years of education rated the 24 familiar fruits and vegetables pairwise on each of the sensory features in separate ratings -- color, shape, taste, touch -- and semantic similarity on a seven-point scale (1: most dissimilar, 7: most similar), with 276 pairs per feature rating. Note that the object sensory features were chosen to form a transition of visual dependence: object color whose qualia is exclusively vision-dependent; object shape whose qualia is dominantly perceived through vision but also could be sensed through touch; object taste and touch whose qualia might be affected by vision but largely determined by nonvisual sensory modalities. Object semantic relationship is often assumed to be abstracted away from sensory experiences. Two blind participants did not complete the touch similarity rating. The ratings were also collected from an independent group of 21 college students (SC2) as a benchmark for between-blind-sighted comparison. For each object feature, group-mean representational similarity matrices (RSM) were obtained by averaging across individual similarity matrices in each group (Fig. 1A; Fig.S2A). Multi-dimensional scaling (MDS;*16*, *17*) using an individual differences scaling solution (INDSCAL) was conducted on the individual RSMs from each group to visualize the object feature space in each group, using the PROXSCAL procedure in SPSS Statistics 19. Two-dimensional object spaces were created for each group. In addition to the object space, the INDSCAL solution also reveals a subject space within which each individual subject is located according to their weights on the dimensions of the identified object space. The weights indicate the importance of each dimension to each individual’s data and the squared distance of each individual subject to the origin of the subject space approximates the proportion of variance of the subject’s data explained by the group solution (Davies and Coxon, 1982). Individual variance within CB and SC group was measured using a leave-one-out procedure, computing the correlation between each individual subject’s RSM and the group-mean RSM excluding the corresponding subject for each group.

#### Object color name generation and color name similarity rating

We also asked the 13 CB and 14 SC participants to generate color names to each of the 24 fruits and vegetables. Participants were encouraged to list as many colors as they considered relevant to each item in the order of relevance. The number of color names produced for each item was not limited. For each item, we listed all color names produced by all the participants from both groups and counted the times that each color name was generated for that item across participants in each group, resulting in a 24 (items) × 9 (color names) matrix for each group (Fig. S1A). We then correlated the items in each group and thus obtained a 24×24 object RSM for each group. Comparison between these RSMs and RSMs obtained through direct object color similarity ratings were carried out to investigate how well the object color space could be approximated by object-color verbal association.

We further approximated the object color space by combining object color name occurrences and the association between color names. We collected 7-point (1: most dissimilar, 7: most similar) pairwise color similarity ratings from both groups to measure the relationship between 9 color concepts (i.e., red, orange, yellow, green, cyan, blue, purple, pink, gold) (Fig. S1B). To combine object color name occurrences with color name associations, we identified, amongst the 9 frequent color words, the most frequently generated color for each fruit/vegetable item in the object color generation task as the representative color for that item. Then, the color similarity between fruit/vegetable items was defined as the similarity between the corresponding representative colors obtained from the color word similarity rating, resulting in a 24×24 object RSM for each group.

### Procedures for task-fMRI experiment

An object color verification fMRI experiment was performed to obtain object color representations in the sighted and congenitally blind people, using spoken names of fruit and vegetables as stimuli. In addition, a color perceptual localizer experiment was conducted on a group of 14 sighted participants to functionally localize brain regions underlying color perception. Seven participants took part in both fMRI experiments.

#### Object color verification experiment

Participants listened to spoken names of 24 fruit and vegetable items and verified whether the item was red or not (Fig. 2A). Participants pressed a button with their right index finger to give a “yes” response and pressed another button with right middle finger for a “no” response. There were three runs. Each run lasted 346s, consisted of 72 3s-long object trials (1s auditory word followed by 2s silence) and 36 3s-long null-trials (3s silence). Each object item was presented three times within each run. The order of the 108 trials was pseudorandomized, with the restriction that no two consecutive object trials were identical and the first and the last trials were object trials rather than null trials. Each run began with a 12s silence period and ended with a 10s silence period. Across the whole experiment, each of the 24 fruits and vegetables was presented 9 times. Independent pseudo-randomizations were created for each run in each participant.

#### Color perceptual localizer

The color perception localizer was an fMRI-adapted version of the Farnsworth-Munsell 100 Hue Task, which has been used previously to identify brain regions involved in color perception (Beauchamp, 1999; Simmons et al., 2007). During the experiment, participants saw blocks of 7 colored or grayscale wheels (Fig.3A) and judged whether the five wedges making up the wheel were uniformly ordered from lightest to darkest hue. Participants were asked to press a button with their right index finger if they found that the hues in the wheel were uniformly sequenced or otherwise press another button with the right middle finger. There were 4 runs in the color perception localizer. Each run lasted 230s, consisted of 3 color wheel blocks and 3 grayscale wheel blocks, with all 6 blocks separated by 15s-long fixation periods. Each block lasted for 21s, consisted of seven 2.5s-long wheels separated by a 0.5s black inter-stimulus interval. The block order was counterbalanced across runs. Each run began with a 12s-long fixation period and ended with a 17s-long fixation period.

### Image Acquisition

All functional and structural MRI data were collected using a Siemens Prisma 3T Scanner with 20-channel head-neck coil at the Center for MRI Research, Peking University.

The functional data were acquired with a simultaneous multislices (SMS) echoplanar imaging sequence supplied by Siemens (slice planes scanned along the rectal gyrus, 64 axial slices, phase encoding direction from posterior to anterior, repetition time (TR) = 2000 ms, echo time (TE) = 30 ms, multi-band factor = 2, flip angle (FA) = 90°, field of view (FOV) = 224 mm × 224 mm, matrix size = 112×112, slice thickness = 2 mm, gap = 0.2 mm, voxel size = 2 × 2 × 2.2 mm). The resting-state scanning lasted 8 minutes (240 volumes), during which the participants were asked to close their eyes and to not fall asleep. In addition, a high-resolution 3D T1-weighted anatomical scan was acquired using the magnetization-prepared rapid acquisition gradient echo (MPRAGE) sequence for anatomical reference (192 sagittal slices, TR = 2530 ms, TE = 2.98 ms, FA = 7°, FOV = 224 mm × 256 mm, matrix size = 224×256, interpolated to 448×512, slice thickness = 1 mm, voxel size = 0.5 × 0.5 × 1 mm).

### Data preprocessing

Task-fMRI data were preprocessed using Statistical Parametric Mapping software (SPM12; http://www.fil.ion.ucl.ac.uk/spm12/). For each individual participant in each experiment, the volumes (n=6) of the silence/fixation block at the very beginning of each functional run were discarded for signal equilibrium. Task-fMRI data first underwent slice timing correction. Then the images from multiple runs were aligned to the individual’s first image of the first run using six rigid body transforming parameters. For multi-variate analyses, the realigned images were not further normalized or spatially-smoothed and the structural images were co-registered to the mean functional image. The deformation fields for the transformations between the Montreal Neurological Institute (MNI) space and the native space were obtained through segmentation. For univariate analyses, the realigned images were further normalized into the MNI space using unified segmentation and spatially smoothed using a 6mm full-width half-maximum (FWHM) Gaussian kernel.

Resting-state fMRI data were preprocessed using SPM12 and the toolbox for Data Processing & Analysis for Brain Imaging (DPABI, v3.1, http://rfmri.org/DPABI)(*31*). For each participant, preprocessing follows conventional procedures, which include: 1) discarding the first 10 volumes for signal equilibrium; 2) slice timing; 3) correction for head movement with rigid body translation and rotation parameters; 4) normalization into MNI space by DARTEL(Ashburner, 2007); 5) spatial smoothing with 4mm FWHM Gaussian kernel; 6) removing the signal trend with time linearly; 7) band-pass (0.01-0.1Hz) filtering; 8) regressing out nuisance variables including the six rigid head motion parameters, the global signal averaged across the whole brain, the white matter signal averaged from the deep cerebral white matter and the cerebrospinal fluid signal averaged from the ventricles to further reduce non-neuronal signal confounds. Note that one blind participant was excluded from the subsequent resting-state functional connectivity analysis due to excessive head motion (> 2mm/2°).

### Data analysis

For each participant, general linear models (GLM) were built to obtain condition-specific beta estimates for subsequent analyses. For the event-related designed object color verification experiment, the unsmoothed, un-normalized data were analyzed using the GLM. The GLM for each run contains 24 regressors corresponding to the 24 fruit and vegetable names. For the block design color perceptual localizer, the smoothed, normalized data were analyzed. The GLM for each run contains two regressors corresponding respectively to the chromatic and achromatic conditions. For both experiments, no participant had head motion larger than 1.5mm/1.5°. Six head motion parameters were further included in each GLM as regressors of no interests to control for potential confounding of head motion. Each regressor was convolved with a canonical HRF and a high-pass filter cut-off was set as 128s for the GLM of each run.

To investigate the object color representation in the brain of CB and SC participants, we conducted Representational Similarity Analysis (RSA)(Kriegeskorte et al., 2008) using a searchlight procedure (Kriegeskorte et al., 2006) in each individual participant. The t-value (each condition relative to baseline) images of the 24 fruit and vegetable names in the object color verification experiment were calculated to capture the activation patterns. The t-images were chosen to suppress the contribution of voxels with high beta estimates due to high noise (Misaki et al., 2010). For each participant, the structural image was co-registered to the mean functional image and segmented into different tissues. The resulting gray matter probabilistic image was resliced to the same spatial resolution as that of the functional image and thresholded at one-third to generate a binary mask for RSA searchlight mapping. For each voxel within the gray matter mask, the multi-condition activation patterns within a sphere (radium=6mm) centered at that voxel were extracted to calculate the neural representational similarity matrix (RSM) based on that search sphere. Pearson correlations were computed between conditions and a 24×24 neural RSM was obtained for each search sphere and then compared with behavior-derived object color RSM using Spearman’s rank correlation, or using partial Spearman’s rank correlations to control for effects of other object features. The resulting correlation maps were Fisher transformed, normalized to the MNI space and spatially smoothed using a 6mm FWHM Gaussian kernel for subsequent group-level statistical analyses.

To test for significant object color representation in both groups, the correlation maps of all CB and SC participants were entered together into a group-level random-effects one-sample t-test analysis using the permutation-based statistical nonparametric method mapping (SnPM; http://go.warwick.ac.uk/tenichols/snpm). SnPM-based one-sample t-test analysis was also conducted for each group respectively. The differences between CB and SC groups were investigated using SnPM-based two-sample t-test analysis. No variance smoothing was used and 10000 permutations were performed.

Group-level random effects analyses were first conducted in anatomically pre-defined left anterior temporal lobe and in functionally defined color-sensitive ventral occipitotemporal cortical (VOTC) regions, respectively. These two regions were given special attention because they have been mostly discussed and suggested as candidate brain substrates for non-sensory, abstract knowledge representation (Binder et al., 2005; Noppeney and Price, 2004; Striem-Amit et al., 2018; Wang et al., 2010) and sensory-based representations (Bannert and Bartels, 2018, 2013; Goldberg et al., 2006; Hsu et al., 2011; Martin, 2016; Simmons et al., 2007). The left ATL (Fig. 2B) was defined as the union of six anterior temporal regions according to the Harvard-Oxford Atlas (probability > 0.2). These regions included the temporal pole, the anterior superior temporal gyrus, the anterior middle temporal gyrus, the anterior inferior temporal gyrus, the anterior temporal fusiform cortex and the anterior parahippocampal gyrus, resulting in 4356 voxels (34,848 mm³). The color-sensitive VOTC (Fig. 3A) was defined as regions showing stronger activation to chromatic stimuli relative to gray-scale stimuli (voxel-level P < 0.05, one-tailed, uncorrected) within the cerebral mask combining lingual (47^#^,48^#^), parahippocampal (39^#^, 40^#^), fusiform (55^#^, 56^#^) and inferior temporal (89^#^, 90^#^) regions in the Automated Anatomical Labeling (AAL) template (Tzourio-Mazoyer et al., 2002), resulting in 6524 voxels (52,192 mm³). Note that for the ROI-based rsFC analyses, a more stringent threshold (voxel level FWE corrected P < 0.05 across voxels in the below-described gray matter mask) was adopted to identify the seed ROI for the color perceptual system. Group-level analyses were further conducted in a gray matter mask to explore object color representation in the whole brain. The gray matter mask was defined as voxels with a probability higher than one-third in the SPM12 gray matter template and within the cerebral regions (1^#^-90^#^) in the AAL template, resulting in 111,493 voxels (891,944 mm³).

For the regions detected from the RSA searchlight mapping, we conducted regions of interest (ROI) analyses to further corroborate our findings. For each ROI, the voxel-wise Fisher-transformed correlation values obtained from RSA searchlight were extracted and averaged across all voxels within the ROI for each participant. Bayesian t-tests implanted in the JASP software (version 0.9.2; https://jasp-stats.org/) were then conducted on the correlation values, with a default Cauchy prior width of r=0.707 for effect size on the alternative hypothesis (H1)(Rouder et al., 2012). For the ROIs identified as showing significant object color representation in both CB and SC groups from the combined-group one-sample t-test, Bayesian independent samples t-test was conducted to test whether there was significant difference between the two groups (H1: CB ≠ SC). For the ROIs identified as significant in one group, Bayesian one-sample t-tests were conducted to test whether the object color representation was significant in the other group (H1: group mean > 0).

An additional RSA analysis was conducted to rule out the potential influence of large individual variability of the explicit pairwise rating derived object color representations within the CB group. Specifically, we excluded three CB individuals whose object color RSMs were most dissimilar to the SC’s group-mean object color RSM from RSA (both behaviorally and neutrally) to match the individual variability of the rating-derived object color RSM between CB and SC groups, resulting in twelve CB participants included in this analysis.

The brain results were projected onto the MNI brain surface using the BrainNet Viewer (Xia et al., 2013) (https://www.nitrc.org/projects/bnv/).

## Acknowledgments

We thank Dr. Yangwen Xu for helpful discussions in experimental design and the help in data collection.

## Author contributions

XW and YB conceived and designed research; XW performed research; XW analyzed data; WM and JG designed imaging acquisition protocol; YB, XW and AC wrote the paper.

---

**Fig. S1.**
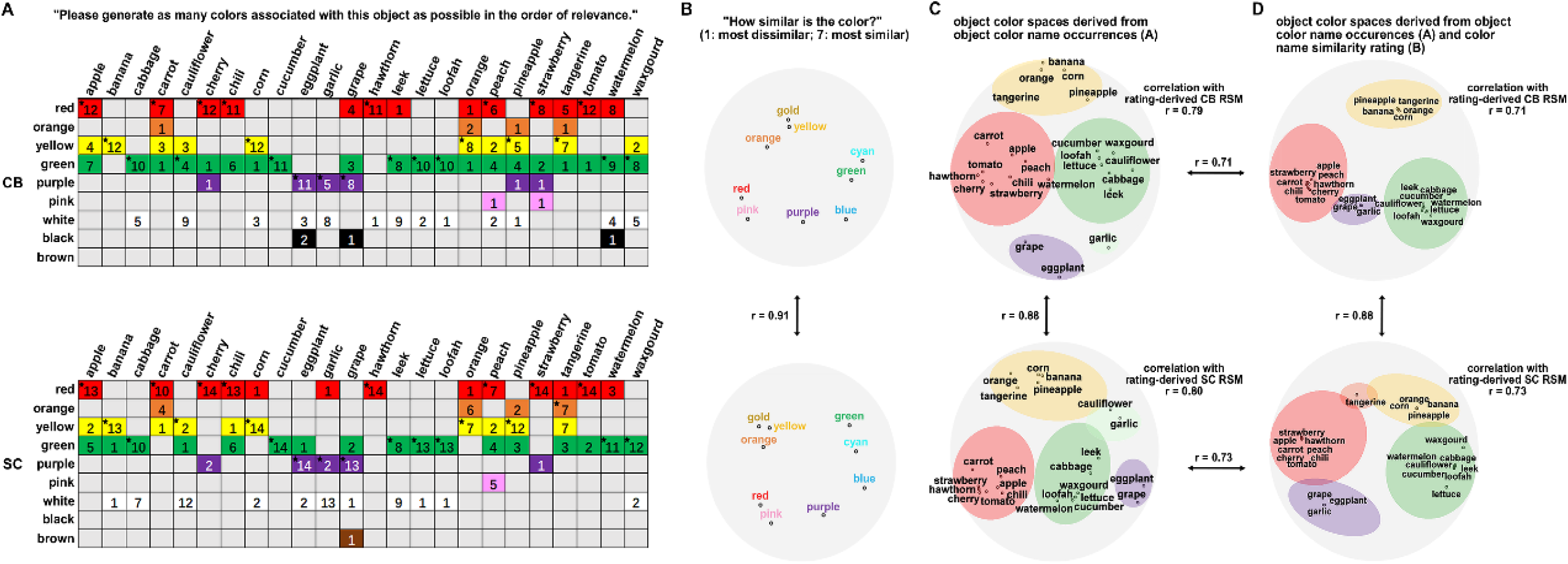
Object color spaces constructed through object color verbal generation and color name similarity rating. (A) Object color name occurrence matrices derived from the object color generation test. The number in each cell indicates the number of times each color was produced for a particular object across all participants in each group. An asterisk denotes the representative color chosen for each object. (B) Color name spaces derived from the color name similarity rating. (C) Object color spaces derived from object color name occurrence matrices (supplementary methods). (D) Object color spaces derived from combining the object color name occurrence matrices with rating-derived color name similarities (supplementary methods). Color patches in (C) and (D) were added subjectively to highlight the clustering pattern of objects with different colors.

**Fig. S2.**
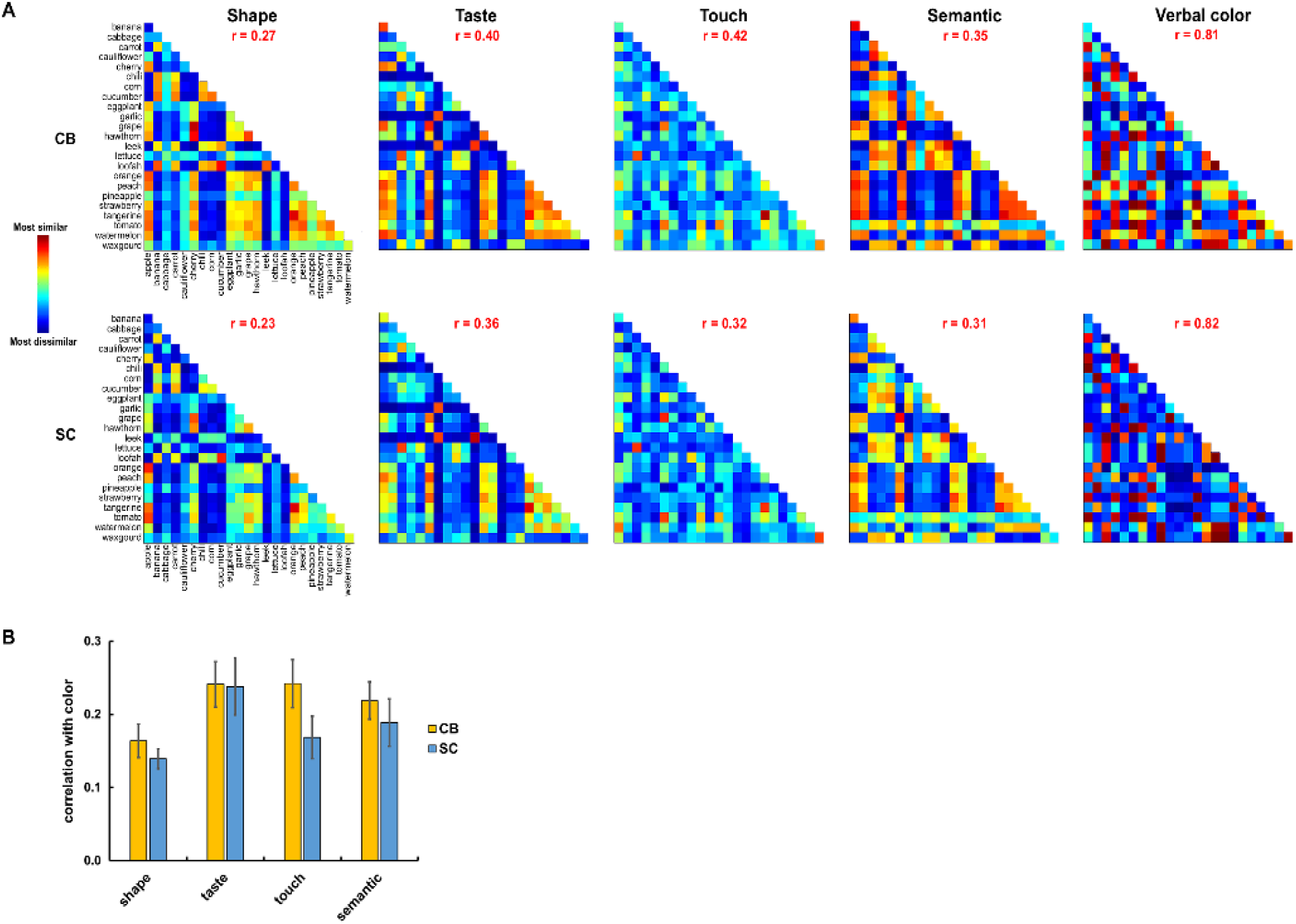
Correlation between object color RSMs and RSMs of other object features. (A) The explicit object feature similarity rating derived RSMs for other object features in CB and SC group. The correlations between the group-mean object color RSM and the group-mean RSMs of other object features are shown (red characters indicate significant correlations). (B) Individual-based correlations between object color RSM and RSMs of other object features.

**Fig. S3.**
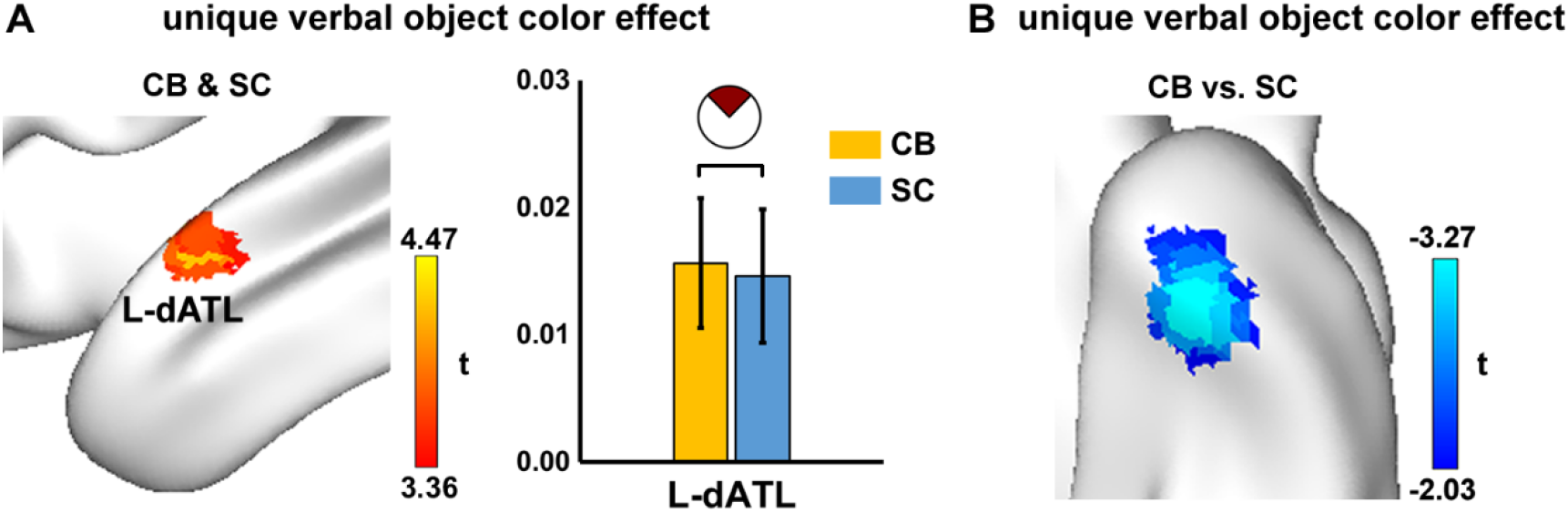
Unique object color representations in CB and SC in the left ATL identified using behavioral RSMs derived from object color name generation and color concept similarity rating. (A) Combination of unique verbal-derived object color effect across CB and SC groups. Voxel-level P < 0.001, one-tailed, FWE corrected P < 0.05, 22 voxels. (B) Between-group comparison of unique verbal-derived object color effect. Voxel-level P < 0.05, two-tailed, 30 voxels. The color bars indicate corresponding t values.

**Fig. S4.**
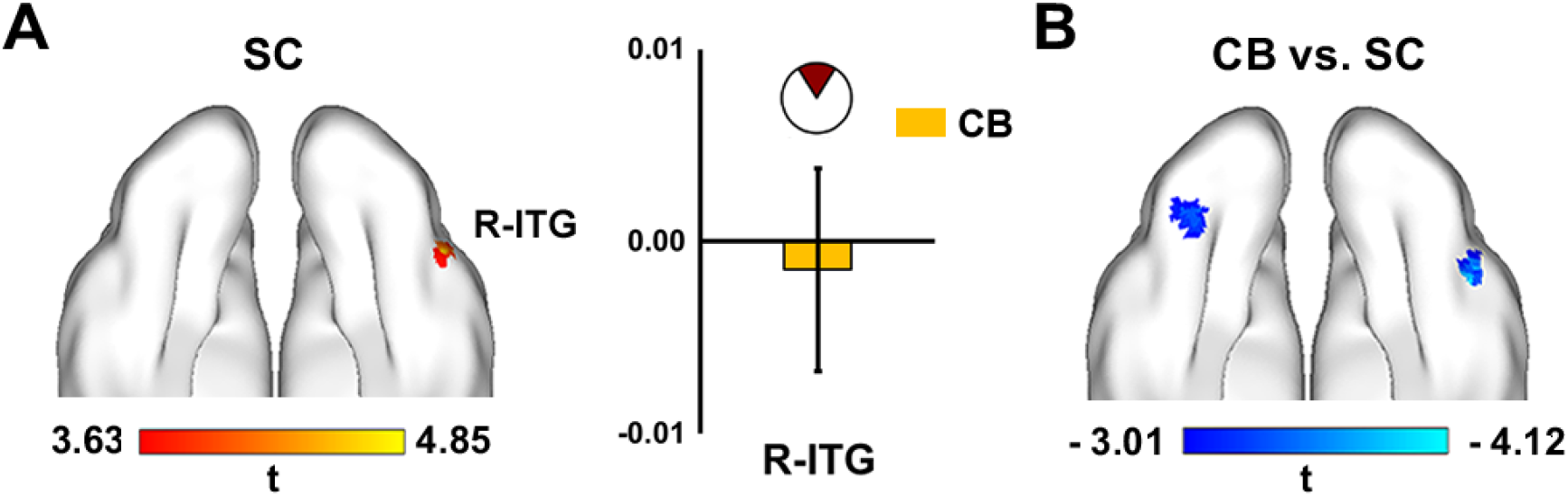
Object color effect without controlling for other object features in the color perceptual mask. (A) Regions showing an object color effect within the color mask in the SC group (voxel-level P < 0.001, one-tailed, cluster size = 15 voxels, uncorrected) and the CB’s object color effect in the identified cluster (bar plot); (B) Regions showing a between-group difference in the color mask (voxel-level P < 0.005, two-tailed, cluster size > 10 voxels, uncorrected). Color bar indicates t values. Negative values indicate weaker effect in CB relative to in SC group.

**Fig. S5.**
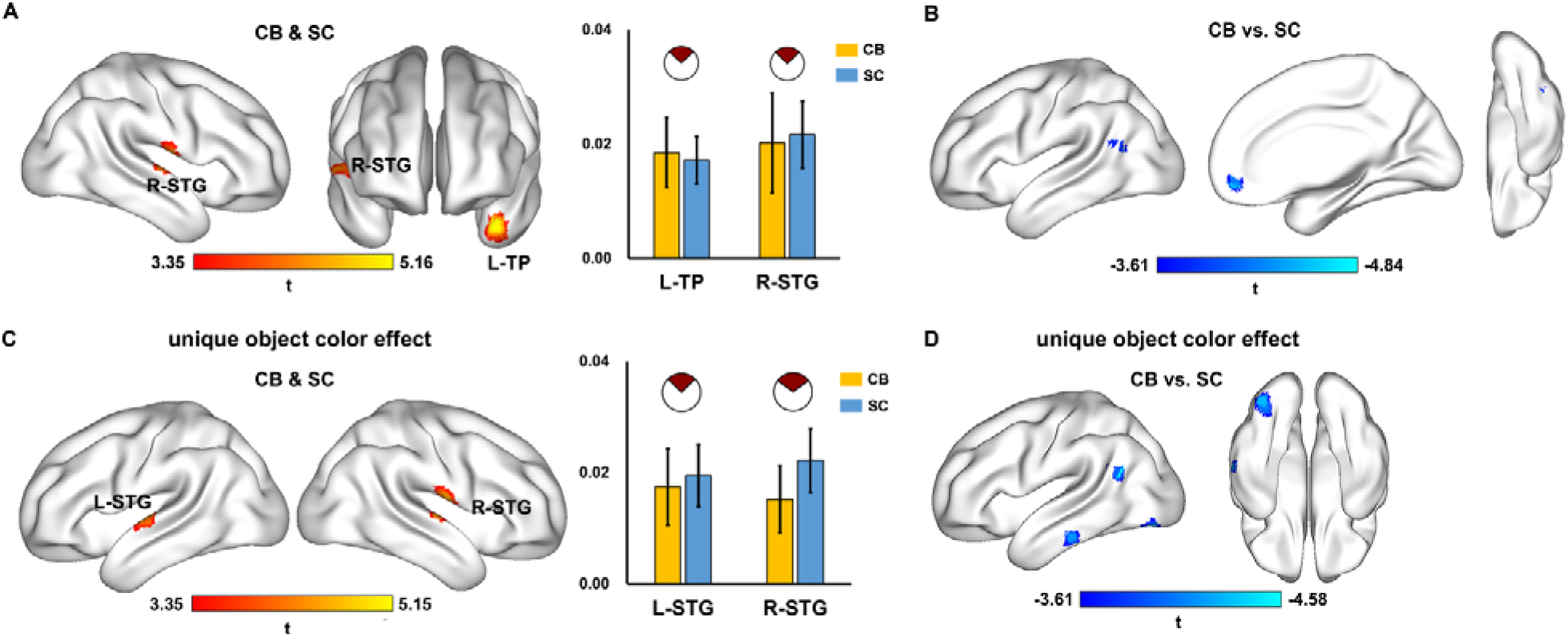
Object color representation in the whole brain. (A) Combination of object color effects across the CB (n=15) and the SC (n=20) groups. Voxel-level p < 0.001, one-tailed, cluster-level FWE corrected P < 0.05. The bar plot shows the extent of object color effect (RSA-derived Fisher-transform r values averaged across participants) in the CB and the SC groups in the identified clusters. The pie chart depicts the odds of the data under H0 (white part) vs. H1 (dark red part). (B) Between-group comparison of object color effect. Cold color indicates weaker effect in CB relative to SC. Voxel-level P < 0.001, two-tailed, cluster size > 10 voxels, uncorrected. (C) Combination of unique object color effects across the CB and the SC groups. The effects of object shape, taste, touch and semantic similarity were removed using partial correlation. Voxel-level p < 0.001, one-tailed, cluster-level FWE corrected P < 0.05. (D) Between-group comparison of unique object color effects. Voxel-level P < 0.001, two-tailed, cluster size > 10 voxels, uncorrected.

**Fig. S6.**
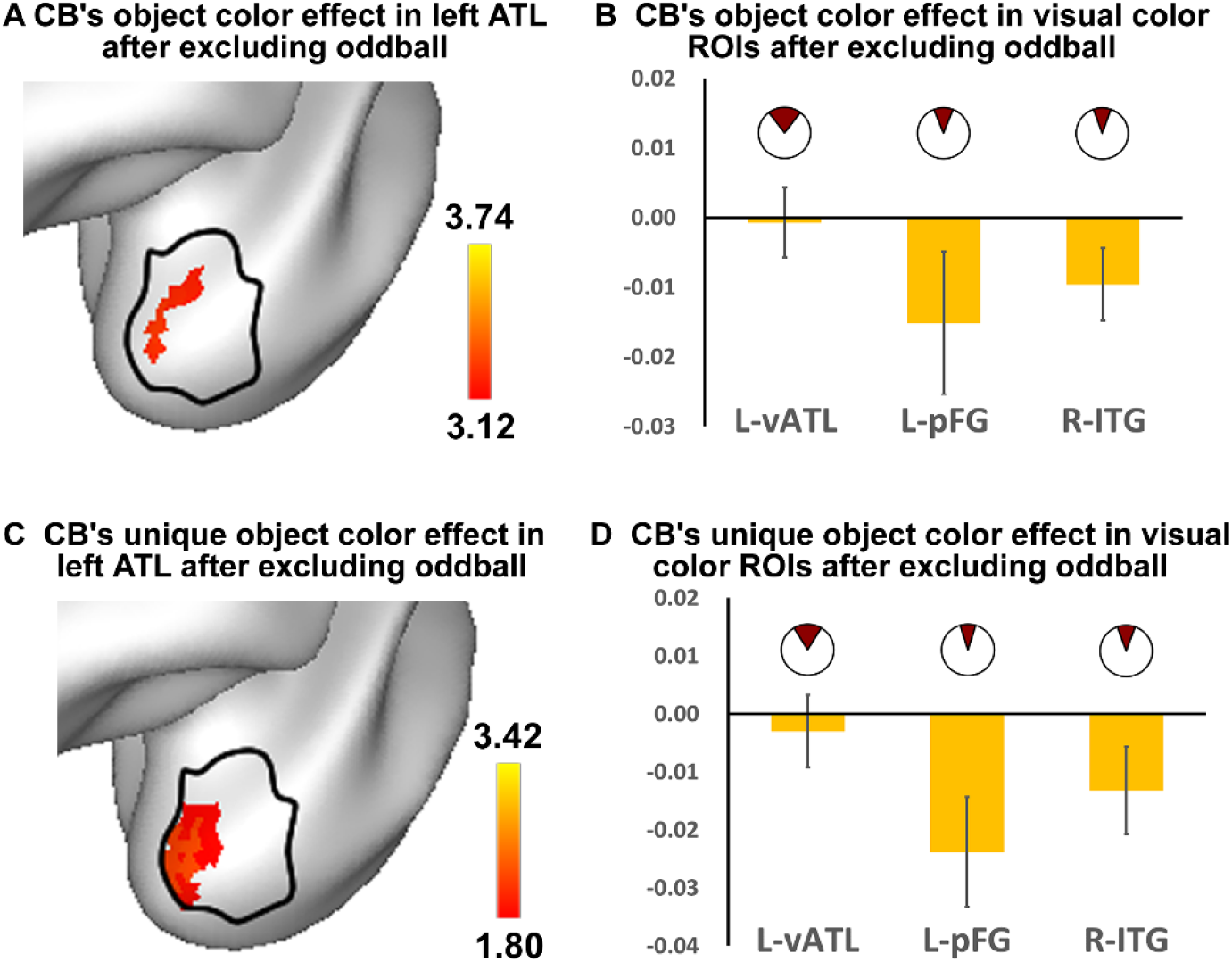
Validation RSA analyses to rule out the potential influence of larger individual variability of the rating-derived object color spaces within the CB group. (A) Regions showing object color effect in the left ATL after excluding three oddball CB individuals whose object color RSMs are most dissimilar to the SC’s group-mean RSM from RSA (both behaviorally and neurally) to match the individual variability of the rating-derived object color RSM between the CB and the SC groups. (B) CB’s object color effect in the visual color ROIs identified in the main text (left vATL from Fig. 2G; left pFG & right ITG in Fig. 3C) after excluding 3 oddball CB individuals. (C) and (D) Unique object color effects (controlling for shape, taste, touch and semantic) in the left ATL and previously identified visual color ROIs after excluding oddball subjects.

**Table S1.** Characteristics of blind participants

| Subject | Gender | Age | Years of education | Handedness | Cause of blindness | Light perception | Experiments participated |
| --- | --- | --- | --- | --- | --- | --- | --- |
| B101 | M | 65 | 9 | R | congenital glaucoma and leukoma | None | All |
| B102 | M | 42 | 12 | R | congenital leukomo | Faint | All |
| B103 | F | 52 | 12 | R | congenital cataracts and eyeball dysplasia | Faint | All |
| B104 | M | 50 | 12 | R | congenital glaucoma | None | All |
| B105 | F | 45 | 12 | R | congenital glaucoma | None | All |
| B106 | M | 50 | 12 | R | congenital microphthalmia, cataracts and leukoma | None | All |
| B107 | M | 59 | 12 | R | congenital eyeball dysplasia | None | All |
| B108 | F | 48 | 9 | R | congenital glaucoma | None | All |
| B109 | M | 35 | 12 | R | congenital microphthalmia and microcornea | None | All |
| B110 | M | 38 | 12 | Bi | congenital microphthalmia | None | All |
| B111 | M | 24 | 15 | R | congenital microphthalmia | None | All |
| B112 | M | 59 | 12 | R | congenital optic nerve atrophy; retinitis pigmentosa; vitreous degeneration | Faint | All |
| B113 | M | 32 | 9 | R | congenital optic nerve atrophy | None | All |
| B114 | M | 36 | 12 | R | congenital eyeball dysplasia due to gestation poisoning | None | task-fMRI |
| B115 | M | 28 | 0 | R | congenital anophthalmia | None | task-fMRI |
| B116 | F | 36 | 12 | R | congenital optic nerve atrophy | Faint | rs-fMRI |

